# Development of a Robust Gel-Free 3D Culture System for Generating Spheroids from Axolotl Blastema Cells

**DOI:** 10.1101/2024.08.29.610223

**Authors:** Zeynep Aladağ Türk, Safiye Serdengeçtі, Sven Vilain, Gürkan Öztürk

## Abstract

Regenerative biology seeks to uncover the principles enabling organisms to restore complex tissues and organs. The axolotl (*Ambystoma mexicanum*), a salamander with unparalleled limb regenerative capacity, remains a premier model system in this field. However, in vitro studies have been limited by the absence of reliable culture platforms for maintaining blastema cell identity and functionality. Here, we present a robust, gel-free 3D culture protocol that enables the formation of spheroids from axolotl blastema cells under fully defined, serum-free conditions. These spheroids preserve the expression of key regenerative markers such as *Prrx1, Msx2*, and *CTGF* for at least 10 days in culture, and exhibit cellular stability supported by antioxidant and amino acid supplementation. Importantly, spheroids cultured for 10 days were capable of initiating extra digit formation when transplanted into limb bud after amputation. With unique advantage of fully defined incubation conditions, this technique adds a powerful in vitro platform to existing experimental models of axolotl limb regeneration.

## INTRODUCTION

Restoration of structural integrity and the function of body parts after damage is the core subject of research in regenerative biology (1-3). An invaluable model organism in this field is the Mexican salamander - axolotl, with unpresented capability to regenerate tissues, organs and appendages (4-12). However, despite extensive in vivo studies on blastema, there is currently no established cell line that retains the essential characteristics of blastema cells for in vitro research. The only known axolotl cell line, AL-1, lacks detailed documentation regarding its origin, characterization, and acquisition methods, which limits its utility in regenerative studies (13). Additionally, existing protocols for primary blastema cultures are plagued by low reproducibility and significant limitations. Kumar et al. (2010) reported that these cultured cells were unsuitable for passaging and could not proliferate(14). Thus, a well-defined and reliable culture technique for regenerative cells of axolotl limb blastema is lacking. In contrast, significant progress has been made in blastema cell culture development in other model organisms like Xenopus laevis, where in vitro cultures have been pivotal for exploring limb regeneration, offering insights into the regenerative capacities and limitations in amphibians (15, 16).

In this study, we present a detailed, robust, and highly applicable gel-free 3D axolotl blastema culture protocol. This system overcomes the limitations of previous methods, allowing the formation of spheroid structures from blastema cells, thus providing a new platform for in vitro studies of axolotl limb regeneration. Our culture system preserves crucial cellular features of blastema, and the spheroids can be harvested to be used in complementary in vivo experiments, paving the way for a deeper understanding of the limb regeneration.

In this study, we present a detailed, robust, and highly applicable gel-free 3D axolotl blastema culture protocol. This system overcomes the limitations of previous methods, allowing the formation of spheroid structures from blastema cells, thus providing a new platform for *in vitro* studies of axolotl limb regeneration. Our culture system preserves crucial cellular features of blastema, and the spheroids can be harvested to be used in complementary in vivo experiments, paving the way for a deeper understanding of the limb regeneration.

### Development of the Protocol

This protocol was developed to overcome the lack of standardized methods for culturing axolotl blastema cells in 3D, which is essential for modeling regeneration in vitro. During method optimization, it was observed that serum-free culture significantly improved spheroid consistency, size, and viability. The use of insulin-transferrin-selenium (ITS), non-essential amino acids, and L-ascorbic acid supported maintenance of blastemal properties for longer periods.

### Applications

This protocol provides a versatile platform for studying regeneration mechanisms in vitro. It can be applied in:

- Functional testing of regenerative cues or factors (e.g., FGF8).
- Genetic manipulation and transgene testing.
- High-throughput drug screening for pro-regenerative compounds.
- Modeling in vitro metamorphosis, which is otherwise technically difficult in vivo due to high mortality rates.

### Limitations

- Long-term maintenance beyond 10 days may alter spheroid phenotype.
- Phenotypic identification of the spheroid cells may be challenging due to their heterogeneous composition and overlapping marker expression patterns.

### Graphical Protocol: Serum-free 3D Culture System for Axolotl Blastema Cells

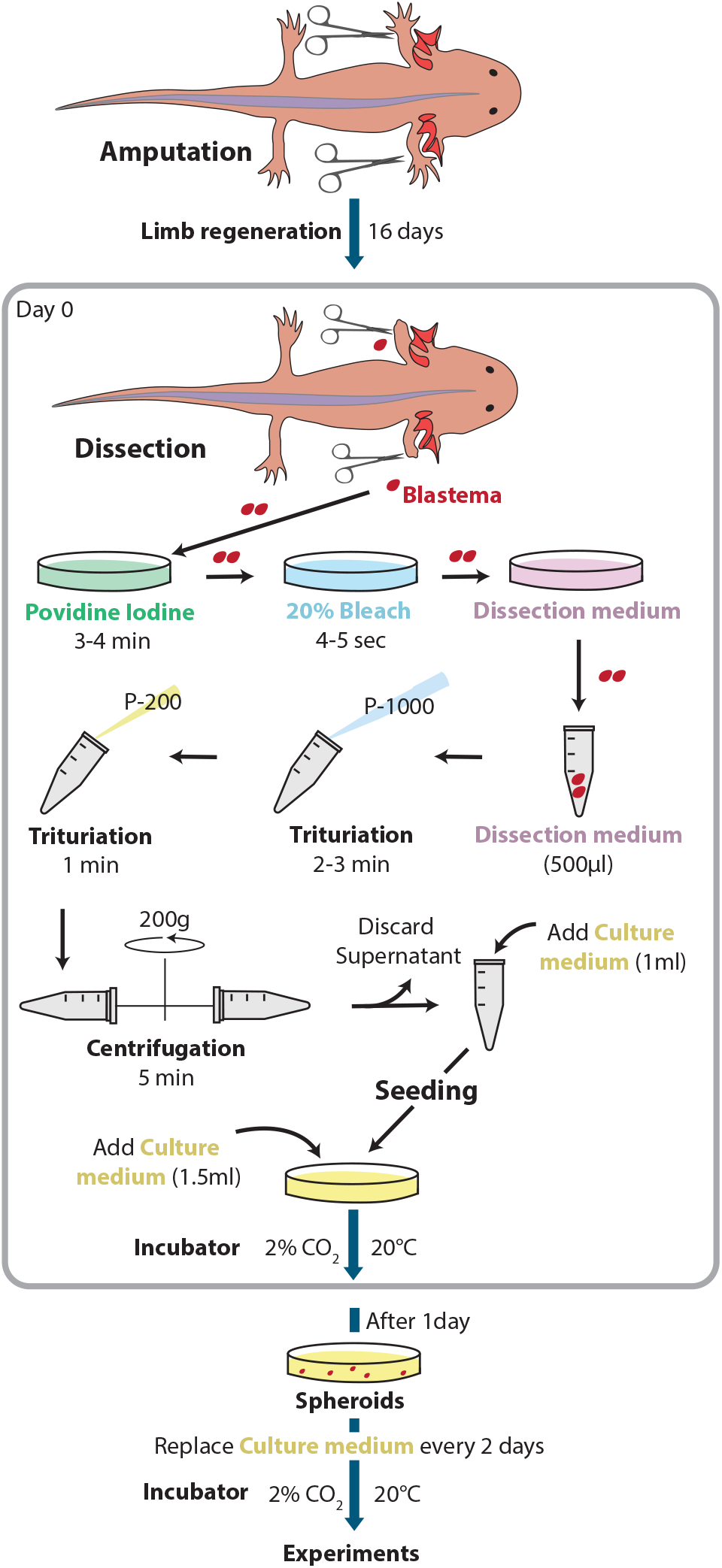

### Schematic overview of the spheroid generation process from axolotl blastema tissue

Day 16 post-amputation blastema tissues are excised, fragmented, and cultured in a defined serum-free medium. Blastema-derived spheroids form in suspension, maintaining regenerative potential and enabling downstream assays such as transplantation and gene expression analysis.

## Materials

- 35 mm Sterile Petri dishes (Nest, Cat. No. 706001)
- 100 cm Sterile Petri dishes (Nest, Cat. No. 704004)
- 2 mL Eppendorf tubes (Nest, Cat. No. 620011)
- 50 ml Sterile Falcon tubes (Falcon, cat. No. 38010)
- Sterile Surgical Design No. 10 Carbon Scalpel Blade
- Micro Scissors
- Iris Scissors
- Pipette Tips (DF100ST, DF200ST, DF10ST)

### Reagents

- DMEM/F12 medium (Multicell, 319-075-CL)
- BSA (Capricorn, BSA-1U)
- FBS (Capricorn, FBS-HI-11A)
- Antibiotic-Antimycotic (Thermo Fisher, 15240062)
- ITS supplement (Thermo Fisher, 41400045)
- Non-essential amino acids (NEAA) (Thermo Fisher, 11140050)
- L-ascorbic acid (Sigma, A0278)
- ddH_2_O
- 1x PBS
- Benzocaine solution (0.3%)
- Povidone iodine Solution
- Bleach (20%)

### Equipment

- AxioZoom V16 fluorescence stereomicroscope
- CT scanner, MILabs U-CT system (for ossification analysis)
- Fluorescence microscope for imaging GFP (LSM800)
- CO_2_ incubator

### Procedure

#### Before You Begin

1. Axolotls (*Ambystoma mexicanum*) measuring 15–20 cm from snout to tail tip were used.
2. Animals were housed under a 12-hour light / 12-hour dark cycle.
3. They were fed every two days.
4. Animals were anesthetized in 0.03% benzocaine solution for 7–10 minutes.
5. After anesthesia, animals were placed on tissue paper to remove excess benzocaine solution.
6. Both forelimbs were amputated midway between the elbow and digits using a Surgical Design No. 10 Carbon Scalpel Blade.
7. Excess skin was pushed back, and protruding bone was trimmed with micro scissors to make the cut area flat.
8. Animals were maintained in tap water under their normal housing conditions for 16 days after amputation.

### Day 0 – Blastema Collection and Dissection

9. On day 16, animals were again anesthetized in 0.03% benzocaine solution for 7–10 minutes.
10. After anesthesia, animals were individually placed on tissue paper to remove excess benzocaine solution.
11. The amputated limb was held proximal to the elbow using curved pattern forceps.
12. Blastemas were excised using 11 cm iris scissors.
13. Harvested blastemas were placed into a 35 mm sterile petri dish containing 1 ml povidone-iodine solution for 3–4 minutes.
14. Blastema tissues were taken out using sterile fine-tip dissecting forceps.
15. They were dipped in 20% bleach solution in a 35 mm sterile petri dish for 4–5 seconds for sterilization.
16. Then transferred into a 35 mm sterile petri dish with 2 ml dissection medium containing:
  - DMEM-F12 (Multicell, 319-075-CL)
  - %1 FBS (Capricorn, FBS-HI-11A)
  - 2% Antibiotic-Antimycotic (Thermo Fisher, 15240062)
  - 10% ddH_2_O

### Preparation of Coated Pipette Tips

17. Pipette tips were coated with coating medium (0.1% BSA in 1x PBS) by aspirating and ejecting the solution to prevent tissue from sticking.
18. Each pipette tip must be coated immediately before use.
19. Coating medium was kept in a 2 ml Eppendorf tube throughout the culture process.

### Blastema Tissue Dissociation

20. Blastema tissues were transferred to a 2 ml Eppendorf tube with 500 µl fresh dissection medium.
21. Tissues were triturated with 1000 µl coated pipette tips for 2–3 minutes.
22. Then triturated again with 200 µl coated pipette tips for 1 minute.

*Note:* The Apical Epithelial Cap (AEC) covering the blastema will remain as tissue fragments and not fully dissociate.

23. The dissociated samples were centrifuged at 200 x g for 5 minutes.
24. Supernatant was discarded.
25. 500 µl culture medium was added to the tube containing the cell pellet. Culture medium composition:
  - DMEM-F12 (Multicell, 319-075-CL)
  - 2% Antibiotic-Antimycotic (Thermo Fisher, 15240062)
  - 10% ddH_2_O
  - 1% ITS (Thermo Fisher, 41400045)
  - 1% non-essential amino acids (Thermo Fisher, 11140050)
  - 10% ddH_2_O
  - 0.2 µM L-ascorbic acid (Sigma, A0278)

### Seeding into Petri Dishes

26. Cells were mixed with culture medium using coated tips.
27. The mixture was added into 35 mm sterile plastic bottom petri dish.

*Note:* Use one 35 mm petri dish per two blastemas to achieve an efficient amount of sphere structures.

28. For larger cultures, multiple blastemas can be pooled and then divided into petri dishes.
29. Each petri dish was filled with 1.5 ml culture medium immediately after seeding.
30. Cultures were incubated in 2% CO_2_ at 20°C.
31. Sphere structures typically begin to appear the next day.
32. Every two days, replace half of the medium with freshly prepared culture medium.
33. This method supports cell viability for up to 1 month.

### Day 3 and Day 10 – Spheroid Collection

34. Pipette tips were coated with coating medium (0.1% BSA in 1x PBS).
35. Spheroids from 3-days-old and 10-days-old cultures were obtained by 1000 µl coated pipette tips into 2 ml Eppendorf tubes.
36. They were centrifuged at 200 x g for 5 minutes and supernatant were discarded.

### Quantitative PCR Analysis

37. Total RNA was extracted from spheroids and blastema samples using TRIGent reagent (equivalent to TRIZol, K5161) following the manufacturer’s protocol.
38. Equal amounts of RNA were reverse transcribed into cDNA using the Advanced cDNA Synthesis Kit (WISENT MULTICELL, 801-100) as per the manufacturer’s instructions.
39. Quantitative PCR (qPCR) analysis was performed using gene-specific primers for Msx2, Prrx1, CTGF (Table 1).
40. The SensiFAST™ SYBR® No-ROX Kit (Bioline-BIO-98005) was used for qPCR reactions.
41. GAPDH was used as the reference gene for normalization.
42. Relative mRNA expression levels were calculated using the delta delta Ct (2^–ΔΔCt) method with negative control.

**Table 1:**
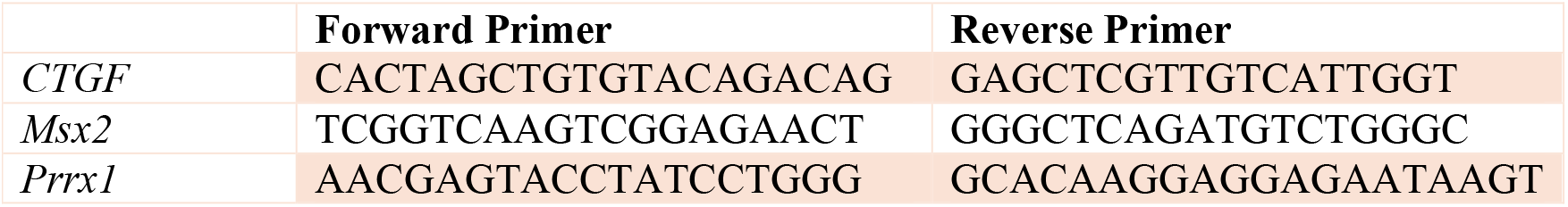
qRT-PCR primers for each gene.

### In vivo Transplantation

43. White axolotls (15–20 cm in length) were anesthetized in 0.03% benzocaine solution for 7–10 minutes.
44. Animals were amputated 16 days prior to transplantation to allow blastema formation.
45. After anesthesia, animals were placed on tissue paper to remove excess benzocaine solution.
46. Animals were then transferred under an AxioZoom V16 microscope.
47. GFP-positive (GFP^+^) spheroids were generated from day-16 blastema tissue of GFP^+^ donor animals, following the protocol described above.
48. Spheroids from 3-day-old and 10-day-old cultures were separately collected into Eppendorf tubes using P20 pipette tips pre-coated to prevent sticking.
49. The collected spheroids were transferred into 35-mm Petri dishes containing minimal medium to maintain viability.
50. GFP^+^ spheroids of varying sizes from 3-day-old and 10-day-old cultures were individually transplanted into the host blastema tissue using coated P20 pipette tips.
51. Each transplantation included 4–5 spheroids of different sizes.
52. Animals were kept moist by covering them with wet tissue paper for approximately 30 minutes after transplantation to allow the spheroid to remain in place within the blastema before being returned to their water tank.
53. Post-transplantation, animals were imaged every two days for approximately six months.

### CT imaging

54. Animals were anesthetized in 0.03% benzocaine solution for 7–10 minutes before each scan.
55. High-resolution micro-CT scans were performed using MILabs U-CT system (MILabs, Netherlands).
56. Scanning parameters: 50 kV, 0.43 mA, 40 ms, voxel size: 80 μm.
57. 3D reconstructions were generated using Recon software (MILabs), and the reconstructions were visualized in Imalytics Preclinical software.
58. Scans were conducted at 55-, 72-, and 148-days post-transplantation.

### Timing Table

| *Step* | *Sub-step* | *Time required* |
| --- | --- | --- |
| **1. Animal preparation and blastema induction** | Amputation surgery (including anesthesia and recovery) | ~30 min per animal |
|  | Post-amputation healing until desired blastema stage (e.g., mid-blastema, day 16) | ~16 days |
| **2. Blastema dissection** | Anesthetizing animal | ~10 min |
|  | Dissection (including transfer to sterile dish) | ~1 min per blastema |
| **3. Tissue dissociation** | Mechanical fragmentation | ~5 min |
|  | Centrifugation | 5 min |
| **4. Spheroid culture setup** | Plating blastema cell/tissue fragments in culture plates | ~5 min |
|  | Incubation under serum-free culture conditions | Variable |
| **6. Optional assays** | qRT-PCR | 2 days |
|  | Spheroid transplantation | ~1 hour |
|  | Micro-CT imaging | ~10 min per animal |

### Troubleshooting Table

| *Step(s)* | *Problem* | *Possible Cause* | *Solution* |
| --- | --- | --- | --- |
| 4, 9 | Animals are not fully anesthetized or wake up early | Low benzocaine concentration, insufficient mixing, or short anesthesia time | Prepare fresh benzocaine solution, keep anesthesia time between 7–10 minutes, confirm unresponsiveness of the animal |
| 13, 15 | Tissue color deteriorates or becomes overly soft | Prolonged exposure to povidone-iodine or bleach | Do not exceed the specified exposure times, use a timer to keep track |
| 20–22 | Tissue does not fully dissociate, large pieces remain | AEC (Apical Epithelial Cap) does not naturally dissociate; insufficient trituration | Do not expect complete dissociation of AEC; small cell clusters are acceptable; carefully extend trituration time |
| 20–22 | Low cell viability | Excessive mechanical stress or using non-coated pipette tips | Always use coating solution (0.1% BSA), perform trituration gently |
| 27–31 | Spheroid formation does not start or is very low | Insufficient cell density or incorrectly prepared culture medium | Use two blastemas per 35 mm petri dish, correctly add and mix culture medium components |
| 32 | Spheroids deteriorate or necrosis begins | Nutrient deficiency or accumulation of metabolic waste | Strictly follow medium change every two days |

## RESULTS

The protocol results in consistent generation of spheroids expressing blastema markers (e.g., PRRX1, MSX2, CTGF). Upon transplantation, 10-day-old spheroids could induce ectopic digit formation.

### Formation of 3D Sphere Structures from Axolotl Blastema Tissues

Cells isolated from day 16 blastema tissue of axolotls were cultured without enzymatic dissociation, resulting in the formation of tissue fragments within the culture. Notably, by the following day, these tissue fragments began to form spherical structures, hereafter referred to as blastemal spheroids (Figure 1A). Over the course of three days, these spheroids progressively got separated from the surrounding tissue fragments and became individual entities (Figure 1B). Furthermore, cells located outside of the tissue fragments were also capable of forming spheroids, though these were significantly smaller in size compared to those derived from tissue fragments (Figure 1C). By using the protocol, spheroids can be obtained from blastema tissue.

**Figure 1:**
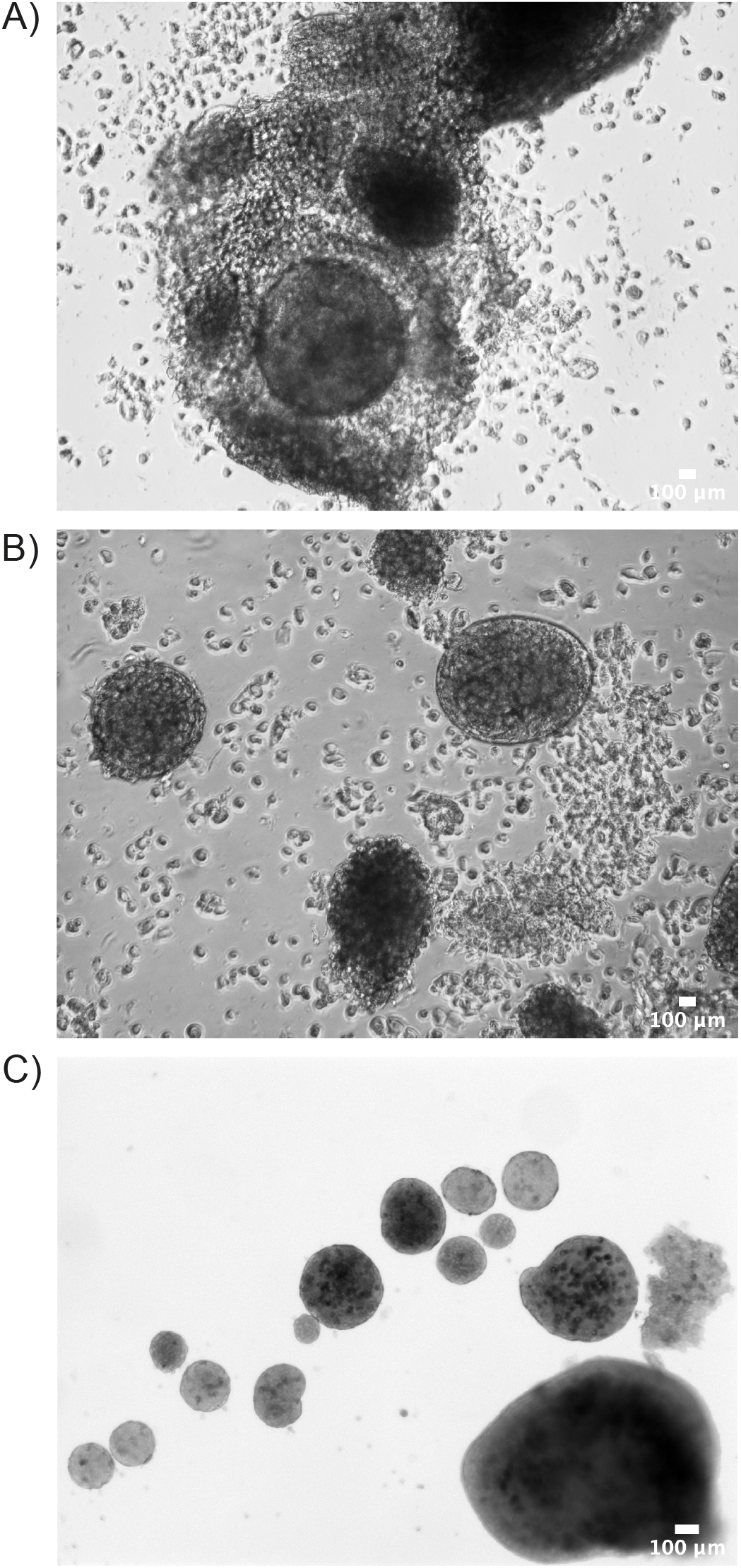
Axolotl blastemal spheroids in culture. Cells were isolated from day 16 blastema of 12 cm animals. A) Spheroids after 24 h in culture; B) Spheroids after 72 h in culture. C) Different sized blastemal spheroids obtained after 10 days in culture. Scale bar: 100 μm.

Furthermore, unlike previously described protocols that induce cell aggregation through mechanical agitation (e.g., rotary shaker systems)(17-19), our method does not require any external force to promote spheroid assembly. Blastema-derived cells spontaneously self-organize into compact spheroids under our defined culture conditions, suggesting that the intrinsic adhesive and signaling properties of these cells are sufficient to drive 3D structure formation in vitro.

### Blastemal Spheroids Retain Expression of Key Blastema Markers

PRRX1 proteins indicate the presence of connective tissue precursors; however, to confirm the blastema-like properties of the spheroids, transcription levels of multiple blastema marker genes needed to be assessed. For this purpose, quantitative reverse transcription PCR (qRT-PCR) analysis was conducted to compare the mRNA levels of key blastema markers, ***CTGF, MSX2***, and ***PRRX1***, between spheroids cultured for 3 days and the original blastema tissue. The analysis revealed that the spheroids expressed comparable levels of these markers to those found in the original blastemas (Figure 2A). Similarly, spheroids cultured for 10 days did not show significant differences in the expression of the same blastema markers compared to the blastema tissue (Figure 2B). Tissue collected from remaining limb immediately after limb amputation was used as a negative control in qRT-PCR normalization calculations. These results suggest that the spheroids maintain their blastemal identity throughout the culture period, as assessed by the expression of key blastema markers.

**Figure 2:**
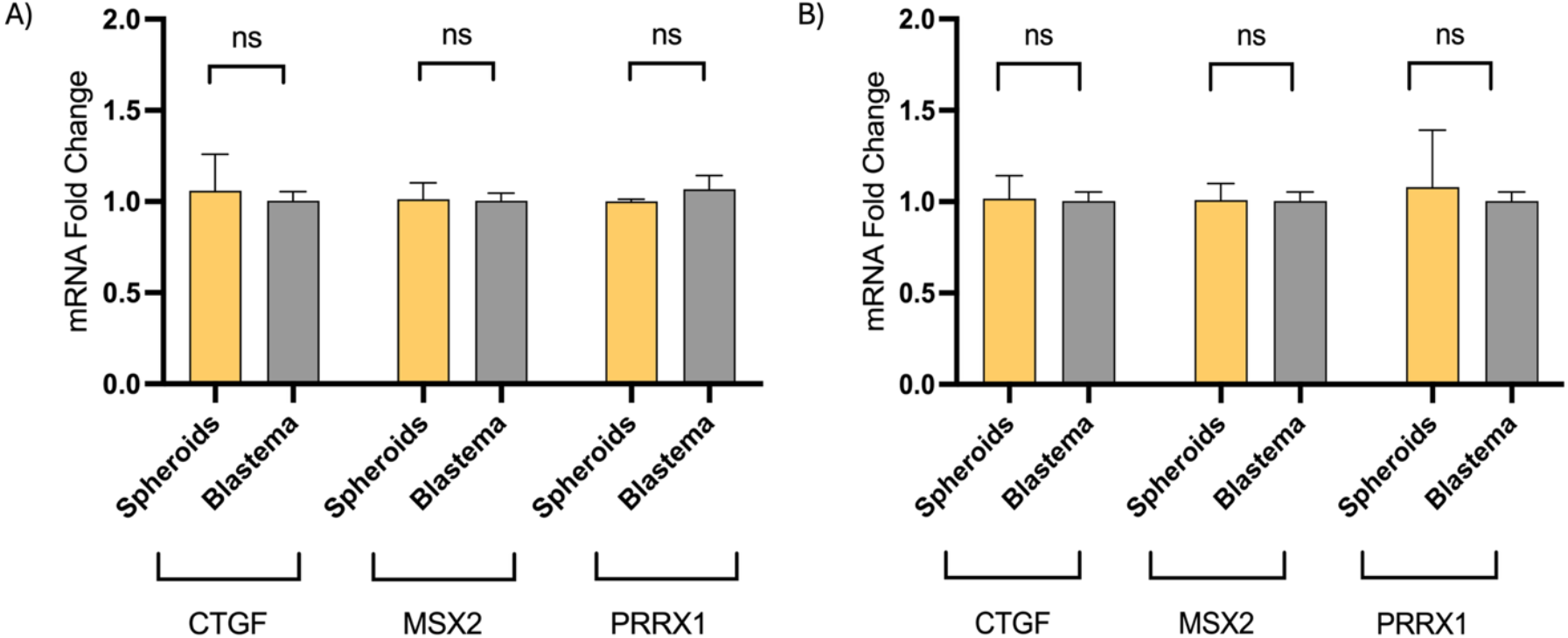
qRT-PCR Analysis of Spheroids and Blastema Tissue. (A) Relative mRNA expression levels of blastema markers in 3-days-old spheroids and corresponding blastema tissue. (B) Relative mRNA expression levels of blastema markers in 10-days-old spheroids and corresponding blastema tissue. Blastema tissues were collected on day 16 post-amputation for both cell culture initiation and direct RNA analysis. No significant differences in marker expression were observed between spheroids and blastema tissue (p > 0.05, Student’s t-test). Tissue collected immediately after limb amputation was used as a negative control in qRT-PCR calculations. n = 6 biological replicates per group. Error bars represent mean ± SEM.

### In Vivo Transplantation Demonstrates Regenerative Potential of Blastema-Derived Spheroids

To evaluate whether the spheroids maintained blastemal characteristics in vivo, we performed transplantation experiments to assess their regenerative potential. Spheroids were generated from GFP+ axolotl blastema tissue harvested at day 16 post-amputation and cultured for either 3 days or 10 days. Following culture, individual spheroids were transplanted into the day 16 blastema tissue of white (non-GFP) host animals to track the fate of the transplanted cells.

In the 3-day culture group, where spheroids were smaller in size, transplanted GFP+ cells dispersed within the host blastema, showing a scattered distribution and integrating into the regenerating tissue without forming a defined structure (Figure 3A).

**Figure 3.**
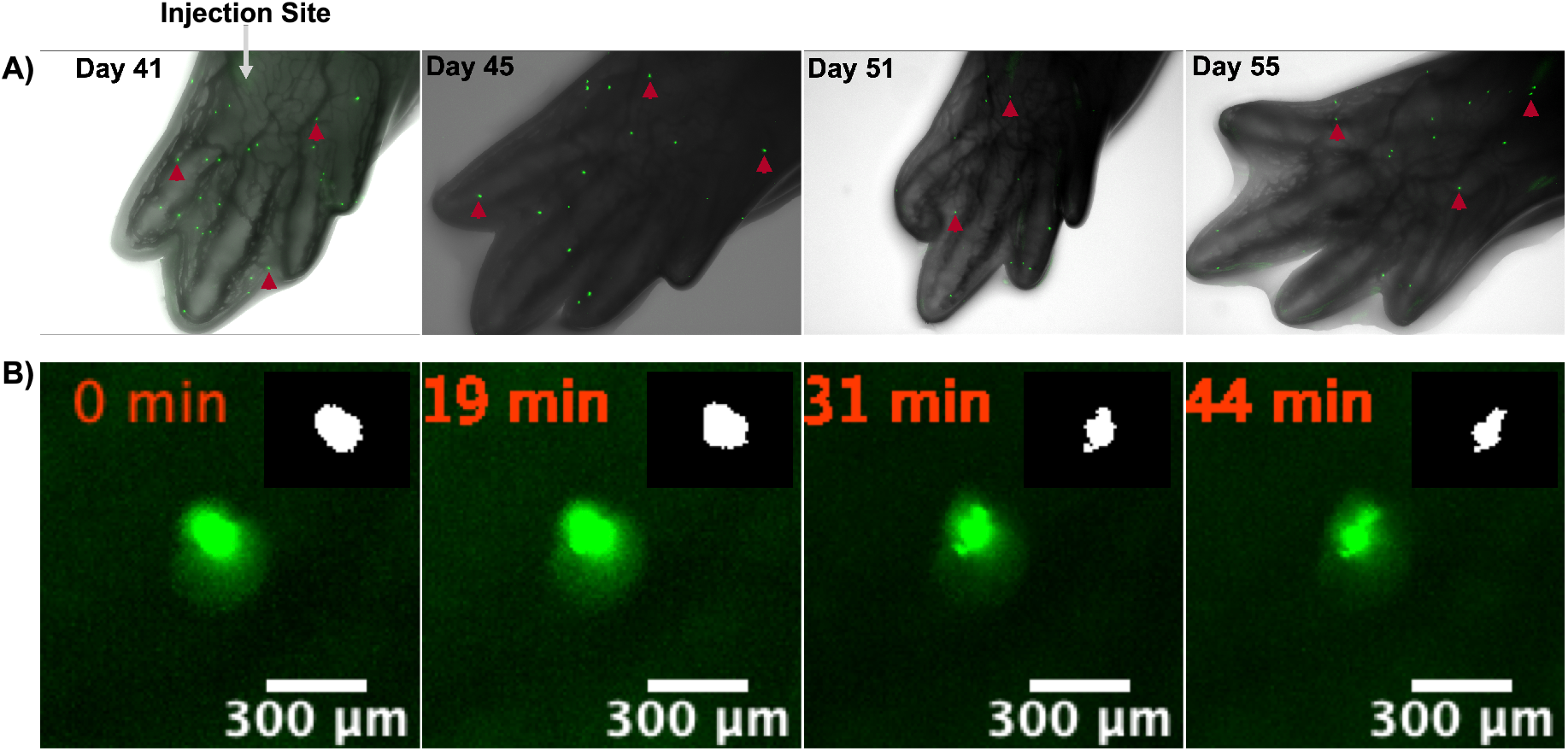
In vivo imaging of transplanted 3-day-old spheroids over time. **Panel A:** Fluorescent imaging of GFP+ cells derived from 3-day-old spheroids transplanted into host blastema tissue, captured at 41-, 45-, 51-, and 55-days post-transplantation. GFP+ cells show a dispersed, scattered pattern within the regenerating limb tissue without forming a defined structure. **Panel B:** Time-lapse imaging of spheroid-derived cells acquired at 1 frame/min for 45 minutes. Four representative frames are shown, illustrating cell movement within the regenerating tissue. Red arrows indicate representative spheroid-derived cells. *Inset shows a binary (black-and-white) magnified view highlighting the morphological changes of the selected cell*.

Time-lapse imaging of transplanted cells from 3-day-old spheroids revealed that GFP^+^ cells extended cellular processes toward surrounding tissue. This behavior suggests potential interactions with host cells and integration into the regenerating environment, despite the absence of a defined digit-like structure (Figure 3B).

In contrast, spheroids cultured for 10 days, which were larger, induced the formation of an additional digit. GFP+ cells were exclusively localized within this newly formed fifth digit and remained detectable in this structure for over 100 days post-transplantation (Figure 4A–C, A′–C′, A″–C″). Furthermore, CT imaging confirmed ossification within the newly formed digit, providing clear evidence that the regenerated digit originated from the transplanted spheroid (Figure 4D-I).

**Figure 4:**
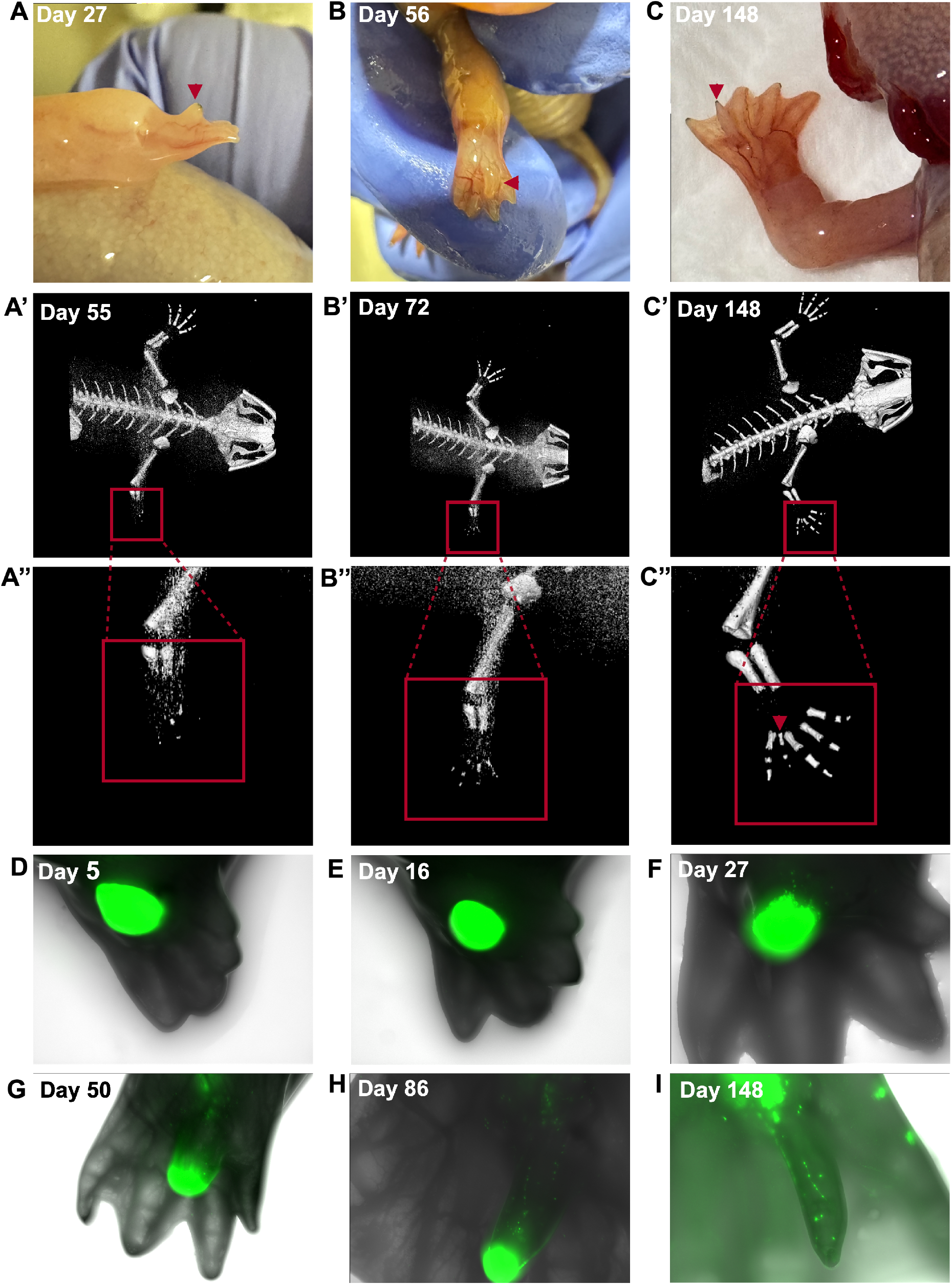
Progressive formation of the ectopic digit after spheroid transplantation. **A–C:** Pictures of growing extra digit over time. **A′–C′:** CT scan images of the transplanted limb at 55 (A′), 72 (B′), and 148 (C′) days post-transplantation, showing progressive bone development within the regenerated digit. **A″–C″:** Enlarged views of the digit region from corresponding CT images (A–C). **D–I:** Fluorescent imaging of GFP+ cells derived from 10-day-old spheroids transplanted into white host blastema tissue, captured at different days post-transplantation. Over time, GFP+ cells were observed to progressively contribute to the generation of an extra digit. By day 148, the new digit was clearly visible formed with both GFP+ and - cells. Red arrows indicate the fifth digit.

## DISCUSSION

This study establishes a robust and reproducible protocol for generating spheroids from axolotl blastema cells under defined, serum-free conditions. Our approach offers a novel in vitro platform to investigate the molecular and cellular dynamics underlying limb regeneration in axolotls. By optimizing the culture environment and confirming the preservation of blastema identity over long periods, this protocol provides a foundational tool to the regenerative biology field.

### Spheroid Formation and Culture Optimization

Axolotl blastema cells cultured on dishes formed compact spheroids within 2 to 3 days after explantation. These spheroids developed from two sources: tissue fragment-based cells and single-cell suspensions, demonstrating that both intact tissue pieces and isolated individual cells were capable of forming spheroid structures in vitro. The heterogeneity in spheroid size and origin may reflect underlying differences in cellular states and regenerative capacity, potentially influencing tissue patterning upon transplantation. During the development of the protocol, we initially tested serum supplementation (%10 Fetal Bovine Serum, Sigma-F4135; data not shown). However, serum negatively affected spheroid formation, leading to irregular morphology and reduced viability, and was thus discontinued. To support spheroid survival and integrity under serum-free conditions, we incorporated supplements including insulin-transferrin-selenium (ITS), non-essential amino acids, and L-ascorbic acid. These supplements were selected based on their known roles in reducing oxidative stress and metabolic burden, thereby enhancing cell viability and maintaining structural integrity. The cultures exhibited compact spheroid morphology with good survival rates under these defined conditions. Detailed descriptions of media composition and supplement concentrations are provided in the Methods section.

An additional advantage of our protocol is that spheroid formation occurs entirely through spontaneous self-organization of blastema-derived cells, without the need for mechanical aggregation systems such as rotary shakers (17-19). This not only simplifies the culture process but also avoids potential shear stress and mechanical artifacts, resulting in spheroids that more closely reflect the in vivo cellular organization during limb regeneration.

### Maintenance of blastema identity in vitro

A key objective of this study was to determine whether spheroids retain their regenerative identity over time. qRT-PCR analysis confirmed sustained expression of key blastema-associated genes, including Prrx1, Msx2, CTGF for at least 10 days in culture. Nonetheless, the consistent transcriptional profile strongly supports the maintenance of a blastema-like state, reinforcing the utility of spheroids as a regenerative model system.

### Functional validation through transplantation and digit morphogenesis

Transplantation of 10-day-old spheroids into the amputation plane of axolotl limbs resulted in the regeneration of a distinct fifth digit. Notably, GFP+ donor-derived cells were exclusively observed within the regenerated digit, with no detectable GFP signal elsewhere in the limb. This spatial restriction strongly supports a direct contribution of the spheroid cells to the regenerated digit, either through structural incorporation or via localized signaling. The fact that the regenerated digit formed precisely at the site of spheroid placement, without broader GFP dissemination, indicates that the spheroids themselves were sufficient to initiate and support digit morphogenesis in a localized and spatially controlled manner.

Importantly, computed tomography (CT) imaging at 148 days post-transplantation revealed the formation of three ossified elements within the regenerated digit, consistent with native skeletal patterning but smaller in size. This anatomical outcome provides strong evidence that the spheroids not only retain regenerative capacity but are capable of supporting both tissue outgrowth and complex skeletal patterning.

Although GFP+ cells were confined to the regenerated digit, they did not appear to proliferate extensively or populate the entire regenerate. This raises the possibility that the spheroids functioned primarily via inductive signaling, rather than forming the regenerate themselves. Such a mechanism would be consistent with blastema-like behavior, where a transient, signaling-competent population instructs host tissue to undergo regeneration. Taken together with sustained expression of blastema-associated genes (e.g., Prrx1, Msx2) prior to transplantation, these results suggest that the spheroids retained a regenerative identity and were sufficient to initiate organized digit morphogenesis, even if their structural contribution was spatially limited.

In an additional transplantation experiment using spheroids cultured for only 3 days, we observed a markedly different regenerative outcome. Instead of producing a consolidated digit, the GFP+ cells became scattered throughout the newly forming tissue without organizing into a defined digit structure. This dispersed integration may reflect the reduced size, maturity, or cellular coordination of the younger spheroid. These findings suggest that the duration of in vitro culture—and by extension, spheroid maturation—may influence the capacity of the transplanted cells to initiate and guide organized digit morphogenesis. Future studies with expanded sample sizes and standardized spheroid metrics will be important to elucidate the relationship between spheroid characteristics and regenerative outcomes.

### Potential applications of the spheroid system

Beyond regenerative assays, the blastema-derived spheroids described in this study offer a unique in vitro platform under defined conditions for modeling axolotl metamorphosis, a process that remains technically challenging to study in vivo due to high mortality rates and systemic effects of hormonal manipulation (20-22). Given their responsiveness, self-organization, and regenerative plasticity, these spheroids could serve as a tractable and scalable model to study cell fate decisions, matrix remodeling, and gene expression dynamics associated with metamorphic transformation. Thus, this protocol may be extended to new experimental domains involving axolotl development, endocrinology, and regenerative transitions. Furthermore, this culture system could serve as a platform for generating blastema-derived organoids or establishing stable blastema cell lines (pending further optimization and analysis), which would provide long-term, renewable resources for studying axolotl regeneration and enable high-throughput screening of regenerative factors.

Regeneration of an amputated limb with proper anatomical alignments and symmetry is an important subject of investigation(23). In the current protocol we use whole blastema tissue to obtain spheroids and they proved to form a complete extra digit when transplanted back to a growing limb bud. Another approach may be selectively recruiting anterior or posterior region cells to investigate their contribution to limb regeneration.

## Conclusion

The establishment of optimized in vitro culture conditions for axolotl blastema cells has enabled the generation of spheroids that maintain key regenerative markers and functional properties. The development of a serum-free, antioxidant-enriched medium has proven essential for spheroid viability and stability. These spheroids not only retain expression of blastema-associated genes such as *Prrx1, Msx2*, and *CTGF*, but also demonstrate regenerative potential when transplanted into injured limbs. Transplantation of 10-day-old spheroids led to the formation of a new digit containing ossified skeletal structures, with GFP+ donor cells contributing specifically to the regenerated tissue. These findings validate the functional relevance of the protocol while highlighting culture duration as a key factor in regenerative success. Furthermore, this protocol holds promise for future applications beyond limb regeneration. In particular, it offers a platform for modeling axolotl metamorphosis in vitro(20, 21). With further optimization and functional validation, these blastema-derived spheroids may serve as valuable tools for dissecting tissue patterning, regenerative signaling, and developmental transitions in a controlled and accessible setting.

## Acknowledgements

We express our gratitude to Dr. Elly Tanaka and her team for generously providing the PRRX1 antibodies. We also appreciate the support from the animal facility at Medipol University, “MEDİTAM,” with special thanks to Ali Şenbahçe for his dedicated animal care.

## Supplementary

**Supplementary Figure 1:**
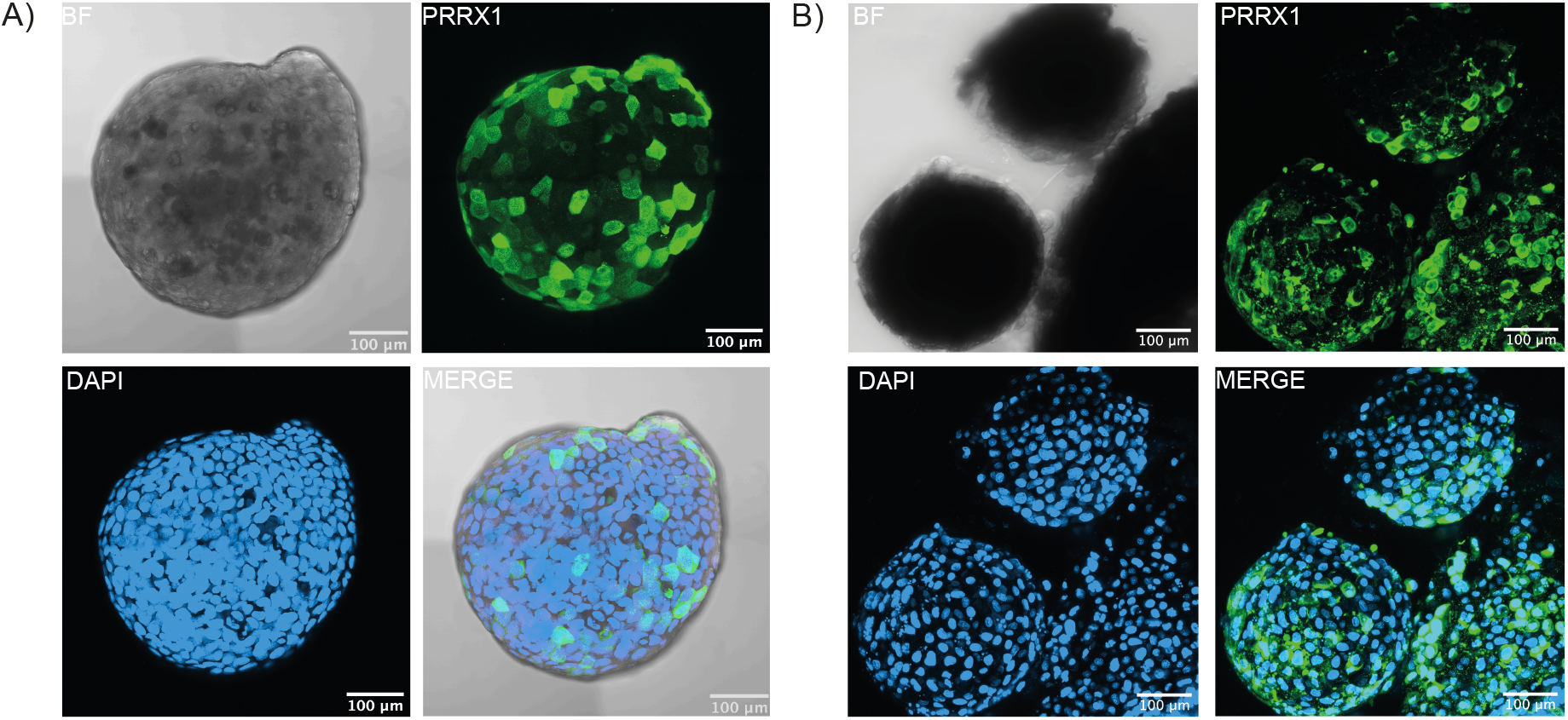
PRRX1 Expression in Axolotl Blastema Cells. Antibody staining of spheroids collected from day 3 and 10 of culture. (A) Expression of PRRX1 in axolotl blastema cells after 72 hours in culture. (B) Expression of PRRX1 after 10 days in culture. BF: Brightfield. n = 3 biological replicates per group. Scale bar: 100 µm.

To assess whether the blastemal spheroids maintain the expression of characteristic blastema markers over time, spheroids were harvested after 3 days (short-term culture) and 10 days (long-term culture). Immunofluorescent staining using a homemade PRRX1 antibody (kindly provided by Dr. Elly Tanaka) was used to detect protein expression within the spheroids. PRRX1 is a transcription factor crucial for controlling gene expression during tissue regeneration. In axolotls, it impacts the formation and differentiation of cells necessary for limb and tissue regrowth. Its role in epithelial-mesenchymal transition (EMT) is essential for the regenerative abilities of axolotl tissues (24-26).

Staining yielded detectable PRRX1 signal in both short- and long-term cultured spheroids (Supplementary Figure 1). However, the observed localization pattern was predominantly cytoplasmic rather than nuclear, which is inconsistent with the known function and intracellular distribution of PRRX1. While the signal may still reflect protein expression --no detection in secondary control staining--the altered localization may be due to antibody performance in axolotl tissue or to technical aspects of our staining procedure. Accordingly, we recommend interpreting these results with caution. To overcome this limitation, PRRX1 expression was also assessed at the transcript level by qRT-PCR, confirming its presence across culture conditions.

**Supplementary Video 1:** Time-lapse video showing spheroid-derived cells captured at 1 frame per minute over a 45-minute period. Imaging was performed using an AxioZoom V16 fluorescence stereomicroscope.

